# Brain network flexibility as a predictor of skilled musical performance

**DOI:** 10.1101/2023.04.26.538360

**Authors:** Kazumasa Uehara, Masaki Yasuhara, Junya Koguchi, Takanori Oku, Sachiko Shiotani, Masanori Morise, Shinichi Furuya

**Author notes:** Corresponding author: Kazumasa Uehara, Ph.D., Associate Professor, Neural Information Dynamics Laboratory, Department of Computer Science and Engineering, Toyohashi University of Technology, 1-1 Hibarigaoka, Tempaku-cho, Toyohashi, Aichi, 441-8580, Japan.

## Abstract

Interactions between the body and the environment are dynamically modulated by upcoming sensory information and motor execution. To adapt to this behavioral state-shift, brain activity must also be flexible and possess a large repertoire of brain networks so as to switch them flexibly. Recently, flexible internal brain communications, i.e., brain network flexibility, have come to be recognized as playing a vital role in integrating various sensorimotor information. Therefore, brain network flexibility may be one of the key factors that define sensorimotor skill. However, little is known about how flexible communications within a brain characterizes inter-individual variation of sensorimotor skill and trial-by-trial variability within individuals. To address this, we recruited highly skilled musical performers (i.e. brass instrumentalists) and used a novel approach that combined multichannel-scalp electroencephalography (EEG) recordings, behavioral measurements of musical performance, and mathematical approaches to extract brain network flexibility.

We found that brain network flexibility immediately before initiating the performance predicted inter-individual differences in the precision of tone timbre (as represented by spectral centroid of the sound), but not trial-by-trial variability at the individual level. Furthermore, brain network flexibility in broader cortical regions, rather than specific local cortical regions, predicted skilled musical performance, indicating that whole-cortical fluctuations determine individual skill. Our results provide novel evidence that brain network flexibility during movement preparation plays an important role in skilled sensorimotor performance and our findings have potentials for designing a new approach to predict an individual’s skill from neural dynamics and a new intervention tool to facilitate physical education.

## Introduction

Our perception, cognition, and motor execution rely on flexible internal brain communications from the microscale to the macroscale level. This communication is well known to be modulated by interactions of the body with the environment (Tognoli and Kelso 2014; Palmigiano et al. 2017). In particular, flexible internal brain communications are indeed necessary to integrate various upcoming sensory information and motor execution because neural information exchanges within the brain vary from minimal to strong, depending on changes in the surrounding environments, requiring the brain to respond to sudden external perturbations. Our brain exhibits spontaneous neural fluctuations that lead to the generation of flexible brain communications and this fluctuation is crucial to brain functioning (Fox and Raichle 2007; Luczak et al. 2009; Anticevic et al. 2012; Bonzano et al. 2015; Cole et al. 2016; Uehara et al. 2022). For example, such spontaneous neural fluctuations and task-evoked brain activity sum linearly during a motor or cognitive task (Fox et al. 2006; Saka et al. 2010; Uehara et al. 2022), suggesting that neural fluctuations partially impact behavior. Therefore, the degree of flexible brain communications is one of the important factors accounting for our cognitive and sensorimotor controls.

In this past decade, brain network flexibility has indeed garnered increasing attention, and several researchers have attempted to quantify this neural trait using blood-oxygen-level dependent (BOLD) signals acquired from functional magnetic resonance imaging (fMRI) (Bassett et al. 2011; Braun et al. 2015; Reddy et al. 2018; Monteiro et al. 2019). These studies extracted the time-varying properties of functional connectivity networks over timescales of seconds to minutes and characterized the network dynamics according to graph-based community structures. This novel analytical pipeline makes it possible to identify the temporal core and periphery of flexible brain regions throughout the whole brain networks. This unique approach can quantify the degree of brain network flexibility as follows: if temporal cores, i.e., brain regional nodes, vary their community assignments over a particular time more frequently, this state indicates more flexible brain networks. On the other hand, if this temporal core maintains consistency, this situation indicates inflexible brain networks (Bassett et al. 2011; Mattar et al. 2016; Bassett and Mattar 2017). Using this neural index, human fMRI studies have sought to investigate how brain network flexibility plays a role in brain functions. It has been reported that brain network flexibility was tightly coupled with motor or cognitive function (Bassett et al. 2011; Betzel et al. 2017). Moreover, it is becoming clear that brain network flexibility is influenced by synaptic transmitters. The excitatory-inhibitory balance, which implies the relative contributions of excitatory and inhibitory synaptic inputs, was found to be involved in brain network flexibility since the A N-methyl-D-aspartate (NMDA) receptor had a potential to modulate the degree of brain network flexibility (Braun et al. 2016). Together, accumulating evidence suggests that brain network flexibility is responsible for orchestrating various brain functions.

Inspired by these pioneering studies, we sought to elucidate a relationship between brain network flexibility and highly-trained skillful motor performance in humans. It is still debatable whether 1) brain network flexibility characterizes inter-individual variation of sensorimotor skill and trial-by-trial variability within individuals, and 2) if so, in what manner brain regions and frequency of neural oscillations in brain network flexibility are associated with these behavioral characteristics. One plausible reason why these open questions have been still unanswered is because previous fMRI studies computed brain network flexibility using BOLD signals but the temporal resolution is sacrificed due to slow hemodynamic response time (typically peaking around 3-5 sec) (Martindale et al. 2003; Yeşilyurt et al. 2008). It indicates that BOLD-based brain network flexibility may be not enough to capture the brain network connections, especially of high temporal neural oscillations. To address these questions, we leveraged scalp electroencephalography (EEG) to record neural signals in a millisecond order, which can provide higher temporal resolution than fMRI. While previous studies computed brain network flexibility during the time window of more than 5-10 minutes of resting state to obtain a large number of data points, our EEG approach allows us to compute brain network flexibility every trial during inter-task events such as the preparation or motor/ cognitive execution. Using this novel approach, we aimed to identify how brain network flexibility can characterize individual differences in the skill and its trial-by-trial variability. For this purpose, we recruited expert musicians who played a wind instrument since they are representative models of skilled individuals and required of preparation for successful performance of skilled motor production. We and others indeed identified behavioral features of their skill while playing a long tone with their own instrument by quantitative analyses of acoustic sounds, which is a proxy for the degree of musical performance skill (Lee et al. 2014; Morris et al. 2018; Uehara et al. 2019).

We postulated that brain network flexibility defined inter-individual differences in skilled musical performance. Here we evaluated precision of tone timbre as an index of musical expertise by calculating spectral centroid of acoustic signals of a tone. Alternative hypothesis that arose from the available evidence was that brain network flexibility throughout the entire brain instead of specific local cortical regions, determined individual musical performance, in accordance with whole-brain network modes (Deco et al. 2009). We assumed that cortical network flexibility is not only determined by activity in individual cortical regions but also emerges as a global property of cortical networks, i.e., large-scale functional organization.

## Methods

Twenty-nine healthy young expert brass instrumentalists (mean age of 24.5 years, standard deviation [SD] of 2.0 years; 8 females) without any history of musculoskeletal and neurological disorders participated in this study. The participants consisted of 20 trumpet, 5 trombone, 3 tuba, and 1 horn player. The average daily practice time was 2.7 ± 1.0 hours. All of the participants had undergone formal musical education at music conservatories and had formal instrumental training. The local ethics committee of Sony approved this study in accordance with the guidelines established in the Declaration of Helsinki. Before starting the data collection, we obtained written informed consent from all participants.

### Experimental task and protocols

Participants sat upright on a chair in front of a monitor, holding own instrument using both hands. To self-evaluate the quality of their own musical performance on a visual analog scale after each trial, participants controlled a foot pedal with their right foot. During each trial, we asked participants to play a visually-guided long tone for 5 seconds without vibrato using their own instruments in response to sequential visual display cues in a dimly lit and soundproof room. We included a simple visual-cued Go/NoGo paradigm in our experimental protocol, as the Go/NoGo paradigm requires participants to withhold a response on NoGo trials (i.e., inhibitory control) and thus characterizes skillful behaviors in musicians (Ruiz et al. 2009). Therefore, we adopted this paradigm in our musical performance task. An experimental environment and trial structure are depicted in Figure 1. After a fixation cross was presented at the center of the monitor for 5 seconds, a “Ready” cue (yellow circle) was displayed for 2 sec with ± 10 % temporal jitter. Next, one of two different colors of cues was presented in a random order within the session. If participants received the visual “Go” cue (green circle), they were asked to play a long tone for 5 seconds as accurately as possible. Recordings of the acoustic sounds started automatically when the initial tone was detected via a microphone. On the other hand, if they received the visual “NoGo” cue (red circle) (Figure 1B), they refrained their musical performance as quickly as possible. Participants rated their performance on a visual rating scale from 1 (worst) to 7 (best) using a foot pedal connected to a PC.

**Figure 1:**
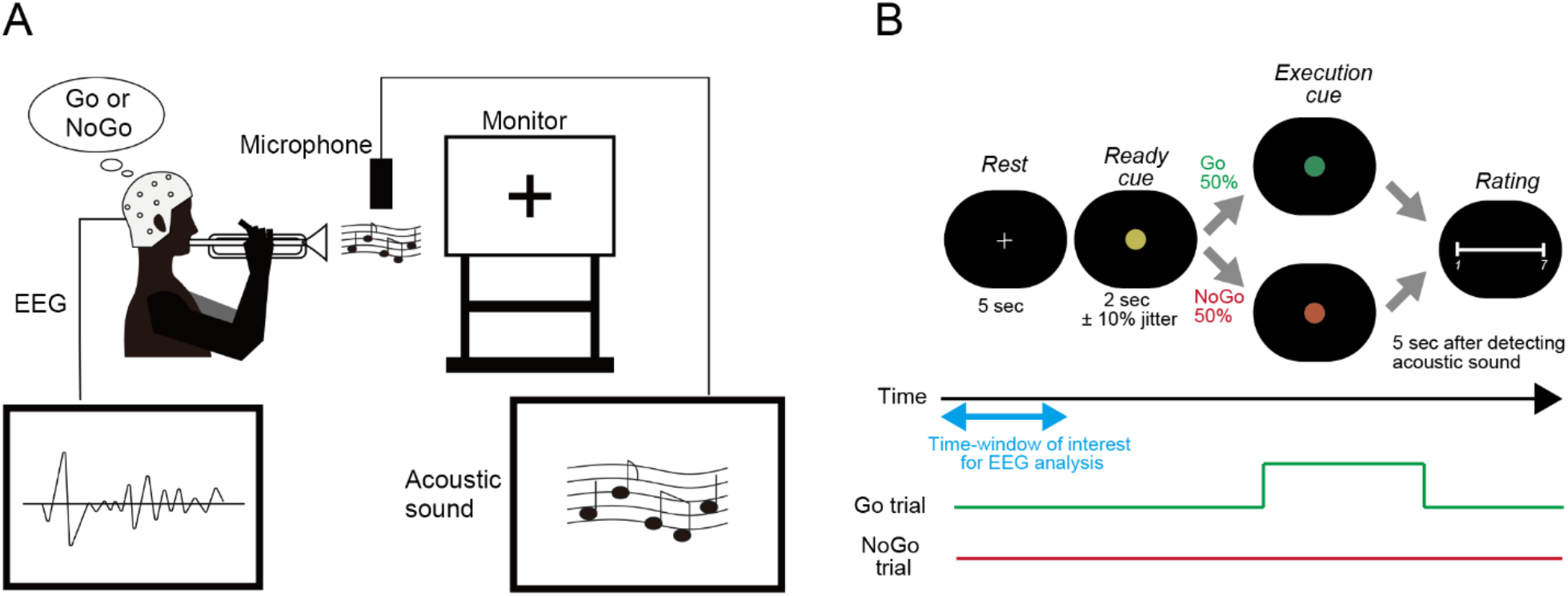
Experimental setup and trial structure. (A) Participants sat upright in a chair in front of a monitor while holding their instruments with both hands. After each trial, they used a foot pedal mouse controlled by their right foot to self-evaluate the quality of their musical performance on a visual analog scale. Acoustic signals of the long-tone sound were recorded using a microphone in a silent environment. Throughout the data collection, 32-channel scalp EEG signals were continuously recorded. (B) Participants were instructed to perform a visually-guided Go/NoGo task while playing a long-tone using their own musical instrument. On each trial, participants were asked to either play a long-tone or cancel their musical performance. After a 5-second rest period and receiving a ready cue (yellow circle) for around 2 seconds, one of two different colored cues was randomly presented within the session. If participants received the visual “Go” cue (green circle), they were asked to play a long-tone for 5 seconds as accurately as possible. On the other hand, if they received the visual “NoGo” cue (red circle), they were asked to inhibit their musical performance as quickly as possible. At the end of each trial, participants rated their performance on a visual rating scale from 1 (worst) to 7 (best) using a foot pedal mouse connected to a PC.

One session consisted of 100 Go and 100 NoGo trials for a total of 200 trials. To secure a sufficient number of trials, we held two sessions in the morning and afternoon (the morning session, started at 10:00 am; the afternoon session, started at 13:30 pm or later) on the same day. Each participant therefore completed 400 trials, which were fed into off-line data analysis. The presentation of the visual cues, onset/offset timing of acoustic signals’ recording, and storage of VAS reports were controlled by open-source software (PsychoPy) implemented in Python. To synchronize the behavior and EEG recordings, a trigger signal was generated by PsychoPy and sent via a digital-to-analog converter (USB-6363, National Instruments, Austin).

### Data recording

#### Acoustic signal

Acoustic signals were recorded with a microphone (C451B, AKG, Vienna) in a silent environment and sampled at 16-bit and 44.1kHz for an off-line analysis. Before starting each session, we checked the recorded sound volumes during the familiarization exercise and made adjustments to avoid signal saturation (i.e. clipping). Recording was automatically initiated when the amplitude of the acoustic signal exceeded 10% of the maximum amplitude.

#### Electroencephalography

During the experimental task, scalp EEG was recorded using a 32-channel electrode cap (WaveGuard EEG cap, Advanced Neuro Technology, Netherlands) according to the 10-10 international system. The ground and system reference electrodes were placed at AFz and CPz, respectively. EEG signals were amplified, digitized with 24-bit resolution and sampled at 1000 Hz using the EEG recording system (eego sports, Advanced Neuro Technology, Netherlands).

Skin/electrode impedance was maintained less than 5 kΩ during data collection. In case a high impedance electrode was found during data collection, the EEG signal was interpolated with the surrounding electrodes throughout off-line analysis using the “*interopl”* function in the EEGLAB toolbox.

### Data analysis

#### Acoustic data

Figure 2 illustrates the analytical procedures for the acoustic data. To quantitatively assess the acoustic sounds, we defined an acoustic feature while playing a long-tone using the spectral centroid. This index is considered as an important feature related to the brightness of timbre and has been proven useful in the analysis of musical instrument sounds (Schubert et al. 2004). The spectral centroid can be defined by the following equation.

**Figure 2:**
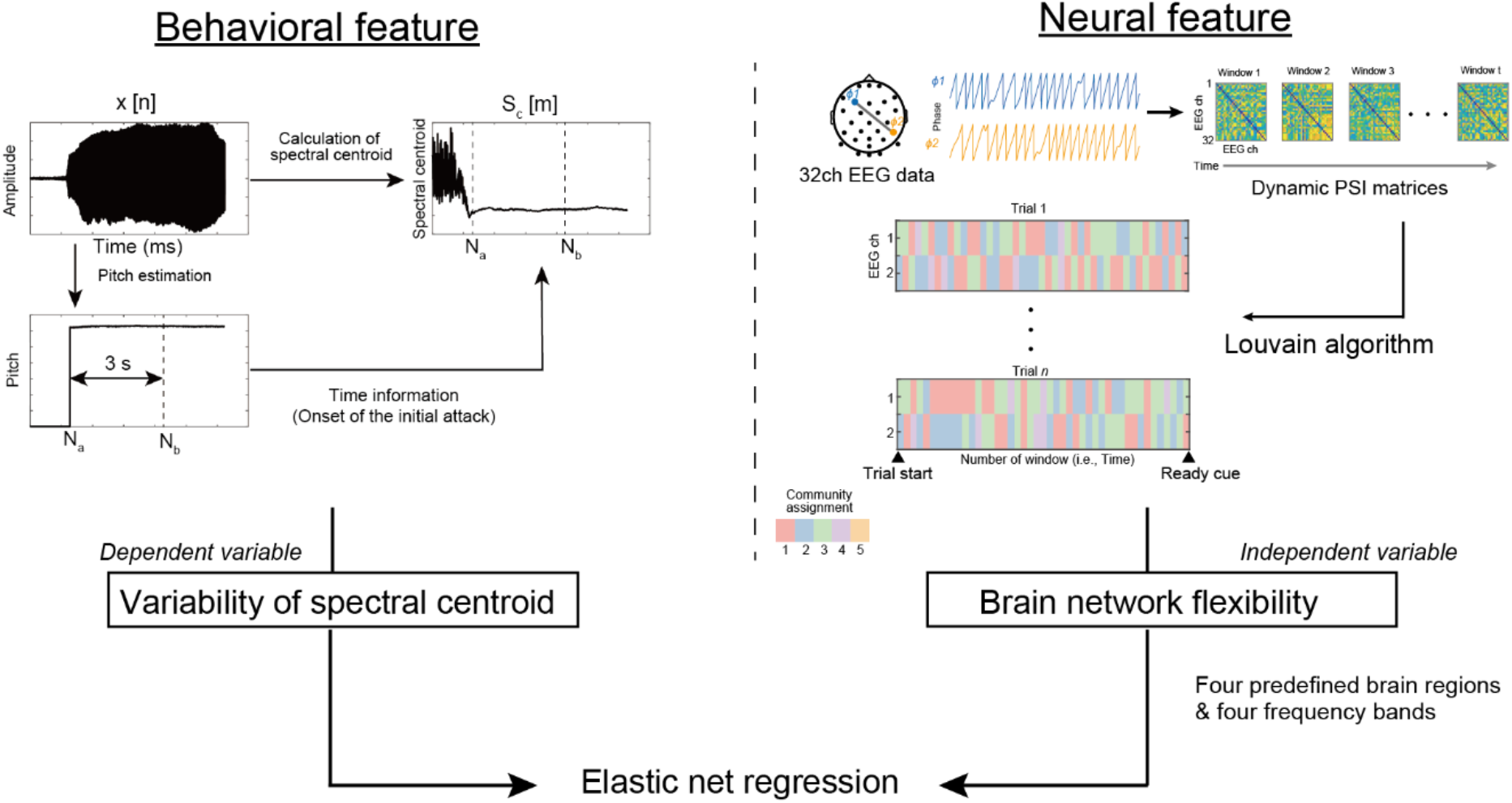
Pipeline for analyzing acoustic and neural data. The spectral centroid, which represents the brightness of timbre while playing a long-tone, was calculated using a spectral-based analysis. The variability of the spectral centroid obtained during a 3 second window was used as a dependent variable. For the neural feature, we first computed dynamic PSI matrices using the phase difference between all channel pairs. The data from these dynamic PSI matrices were fed into a community detection analysis using the Louvain algorithm. The indices of brain network flexibility in four predefined brain regions (frontal, temporal, parietal, and occipital regions) and four frequency bands (theta, alpha, beta1, and beta2) were treated as independent variables. To reveal the relationship between behavioral and neural features, a model of Elastic Net regression was used.

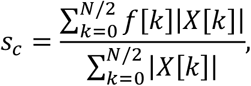

where, *X*[*k*] is the spectrum calculated from the acoustic signal *x*[*n*], and *k* represents the discrete frequency index. *N* is the length of the Fast Fourier Transform, and *f*[*k*] represents the frequency related to the *k*-th index of the discrete spectrum. The spectral centroid is typically defined with respect to the spectrum. To analyze the time-sequence of the spectral centroid, we calculated the spectral centroid *s*_*c*_[*m*] for each frame of the spectrogram using the following equation.

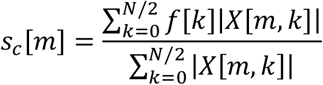

where, *X*[*m, k*] represents the spectrogram obtained from the acoustic signal *x*[*n*], and *m* denotes the frame number. In this study, the analysis was carried out with a frame shift of 5 ms.

We calculated the spectrogram using CheapTrick (Morise 2015), which is a method for estimating the spectral envelope of speech signals, instead of the short-term Fourier transform (STFT). This is because the spectrogram calculated with STFT includes the influence of harmonic structures and negatively affects the time-sequence of the spectral centroid as a pitch-dependent artifact. On the other hand, CheapTrick has the advantage of removing this artifact from the short-time spectrum. This novel approach allows us to obtain the timbre feature without any pitch-dependent effects.

Finally, we calculated the standard deviation of the time-series of the spectral centroid during seconds immediately after the onset on every trial. To identify the first attack of the blowing, we used a fundamental frequency (F0) signal.

In this study, we developed an automatic error detection algorithm to eliminate erroneously recorded trials. This algorithm can detect three types of errors: flying, silence, and redoing. Flying errors occurred when the attack of a note was lost due to sudden initiation of recording, whereas silence errors were observed when there was no performance signal at all. Redoing errors occurred when multiple notes were detected because some participants re-blows in the middle of the performance. As mentioned above, this study used the automatic sound recording system according to the predefined threshold (see, Methods section). First, to deal with background noise in the experimental room, we used a pre-recorded background noise and performed spectral-gating-based denoising (Sainburg et al. 2020). Next, voice and silence flags were calculated for each waveform frame obtained by the STFT. F0 was estimated using the fast DIO algorithm (Morise et al. 2009) in the range from 50 to 2000 Hz. F0 was then re-estimated using a more accurate Harvest algorithm (Morise 2017) in the narrower range from 3/4 to 4/3 Hz of the F0 mode. The frames with the presence of F0 were flagged as voiced. On the other hand, frames with the absence of F0 or the sound pressure level (SPL) were more than 30 dB below the averaged level were considered silent (i.e., correct trial). The range of F0 and threshold of SPL were determined by tuning in some error trials within individuals. The ratio of correct trials is reported in the Results section.

#### EEG data

We performed a series of EEG analyses using functions in the EEGLAB toolbox (Delorme and Makeig 2004) in combination with custom-written MATLAB codes (MathWorks, Natick, USA). Figure 2 illustrates the pipeline of brain network flexibility. Regarding preprocessing, the contentious EEG signal was re-referenced to a common average signal across all channels and then segmented into a 4.5 sec-epoch before the “Go” cue. In this study, we focused on the preparation period immediately before the onset of the musical performance (i.e., long-tone playing) in each trial. This is because we presumed that our brain has to orchestrate the neural network dynamically and flexibly prior to the movement onset to produce skilled performance. Indeed, action preparation is a more dynamic period than one that has been previously considered. For example, the selective suppression of local brain circuits in the motor cortex occurs before forthcoming movements (Hasegawa et al. 2017). Neural preparatory activity, that is not directly coupled to movement output, facilitates motor learning memories (Sun et al. 2022). In a neurophysiological study of musicians, it was reported that predictable auditory information before movements can activate motor representations in an anticipatory muscle-specific manner in order to make sound-related actions (Stephan et al. 2018). These findings allow us to posit that neural preparatory activity may provide the initial state of the dynamical system within the brain that generates the activity patterns of forthcoming movements. We therefore focused on the preparation period to analyze EEG data.

The epoched EEG signals were band-pass filtered between 1 and 50 Hz and 50Hz power line noise was also removed. To remove physiological-related noise, we applied independent component analysis (ICA). Noise-related signals were identified by using visual inspection based on previous literature (Romero et al. 2008; Radüntz et al. 2015) and were then removed from the epoched EEG signals. After ICA-based denoising, epochs containing residual artifacts were rejected from the dataset using an amplitude criterion (±100μV). Finally, we applied the current source density (CSD) transformation using the CSD toolbox (version 1.1) (Perrin et al. 1989; Kayser and Tenke 2006) to attenuate volume conduction effects.

CSD-transformed EEG time series data were decomposed into their time-frequency representations of instantaneous phase with frequency ranging from 1 to 50Hz in 1-Hz steps through a continuous wavelet transformation based on the Morlet wavelet with the number of cycles set to 5 (Lachaux et al. 1999, 2000).

To quantify EEG-based large-scale brain network dynamics, the instantaneous phase synchrony index (PSI) at each frequency was calculated for all possible EEG electrode pairs (total of 496 pairs).

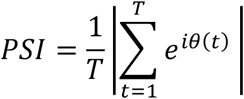

where *T* is the number of time-points that correspond to three cycles of the center frequency with 90% overlapping windows. *i* denotes the imaginary unit, and *θ*(*t*) =φ_m_(*t*)-φ_n_(*t*), in which φ_m_(*t*) and φ_n_(*t*) _represent_ the instantaneous phase of the *m*_th_ and *n*_th_ electrodes at the time point *t*. This index can be considered as the phase relation between two EEG signals from different electrodes at each time point and ranges from 0 to 1, where a value near 0 and 1 implies completely random and perfect synchrony in EEG signals between two electrodes, respectively. The instantaneous PSIs were obtained for every time window and frequency in 1-Hz steps and averaged within the following 4 frequency bands: theta (4-7Hz), alpha (8-12Hz), beta1 (13-25Hz) and beta2(26-40Hz) for each single trial.

To assess brain network flexibility using instantaneous PSI dataset, we used a Louvain-like algorithm (Blondel et al. 2008; Mucha et al. 2010), which is a well-known mathematical method for extracting the community structure inherent in complex networks and has recently been adopted to neuroimaging data such as fMRI and EEG. First, multilayer community detection was performed using the modularity quality function through the generalized Louvain algorithm.

This algorithm can be defined by the following equation:

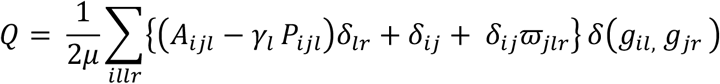

where Q is the multilayer modularity index. *l* implies the number of layers in the multilayer network, μ is the sum of all the edge weights in the network, A_ijl_ is time-by-time N x N functional connectivity matrices (N being the number of EEG electrodes) based on the PSI values, the element *P*_ijl_ is expected weight of the edge between node *i* and *j* under some null model matrix, *γ*_l_ is the structural resolution parameter, which defines the weight of intralayer connections. In this study this parameter was set to 1. *g*_il_ implies the community assignment of node *i* in layer *l, g*_*jr*_ is the community assignment of node *j* in layer *r. ω*_jlr_ is the connection strength between nodes in two layers. δ(*g*_il_,*g*_*jr*_)refers to the Kronecker delta function which equals 1 if g_il_=g_jr_ (i.e., node *i* and node *j* belong to the same module) and 0 otherwise.

Finally, we defined flexibility *f*_*i*_ as the number of times a node changed its module assignment over time, normalized by the total number of possible changes. We then averaged *f*_*l*_ within predefined four brain regions (Frontal, temporal, parietal, and occipital brain regions) per trial using the following equation:

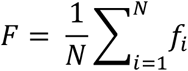

According to human neuroanatomy, four predefined brain regions of interest include the following: the frontal region (8 channels: Fp1, Fp2, F3, F4, Fz, F7, and F8), the temporal region (6 channels: T7, T8, CP5, CP6, P7, and P8), the parietal region (12 channels: FC1, FC2, FC5, FC6, C3, C4, Cz, CP1, CP2, P3, Pz, and P4) and the occipital region (4 channels: POz, O1, O2, and Oz) (Figure 3A).

**Figure 3:**
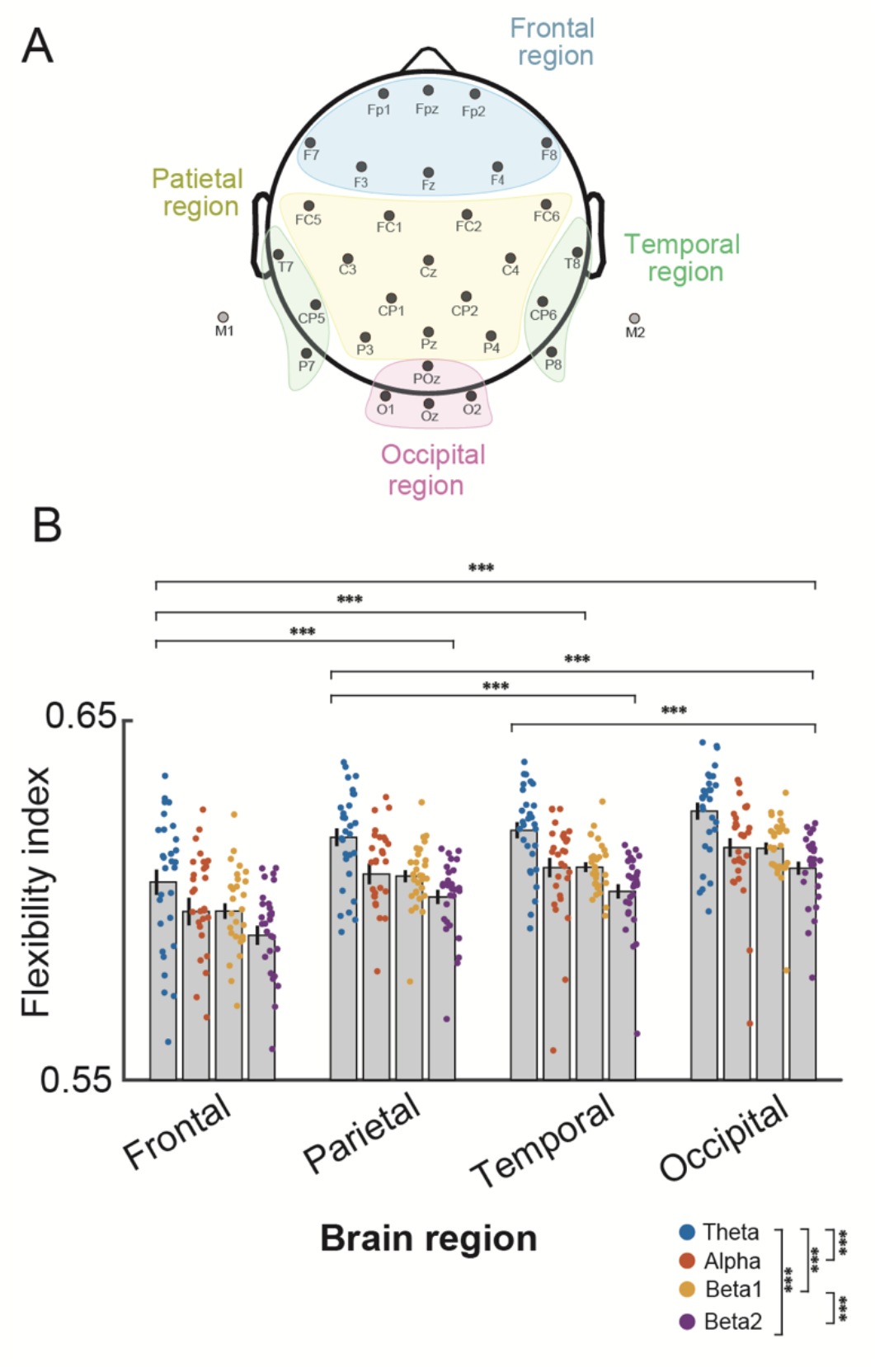
An anatomically-based parcellation of the EEG channels and results of brain network flexibility across each brain region and frequency band during the preparation period. (A) Thirty-two channels, excluding the two mastoid channels (M1 and M2), were classified based on the neuroanatomical locations. Four predefined brain regions of interest were classified: the frontal region (8 channels: Fp1, Fp2, F3, F4, Fz, F7, and F8), the temporal region (6 channels: T7, T8, CP5, CP6, P7, and P8), the parietal region (12 channels: FC1, FC2, FC5, FC6, C3, C4, Cz, CP1, CP2, P3, Pz, and P4), and the occipital region (4 channels: POz, O1, O2, and Oz). (B) Each grey bar plot shows the group-averaged flexibility indices for each brain region and frequency band. The y-axis represents the level of flexibility in brain networks, with high values indicating greater flexibility. Each dot represents an individual, and each error bar indicates the standard error of means across individuals. The statistical significance is indicated by *** p < 0.0001.

#### Statistics

Statistical analyses were performed with the R code (R Development Core Team version 4.1.2, www.r-project.org/). To characterize relationships in the network flexibility (*F*) between each brain region and frequency, first, flexibility indices in each trial were averaged across trials in frequency bands and individuals. We then applied a two-way repeated measures analysis of variance (ANOVA) with Frequency-bands (4 levels: theta, alpha, beta1, and beta2) and Brain regions (4 levels: frontal, temporal, parietal, and occipital regions) as within-subject factors for the flexibility indices. Post-hoc comparisons for ANOVA were applied using Holland-Copenhaver comparisons whenever the main factor was significant (Holland and Di Ponzio Copenhaver 1988). Note that the assumption of sphericity in our repeated measures ANOVA was violated, the p-value was computed with a Greenhouse-Geisser correction.

In this study, the most important issue was the extent to which the flexibility of the cortical network is associated with the superiority of a musician’s performance skills. Here we developed a statistical model that explained the skill of musical performance by brain network flexibility. To test this, we developed an Elastic Net regression model, which takes into account a penalized regression (Zou and Hastie 2005). Elastic net regression allows for selecting variables with a weighted combination of L1 and L2 regularization. Sixteen flexibility indices (4 brain regions x 4 frequency bands) were treated as independent variables. The spectral centroid of the acoustic sound, which indicates musical performance skill, was treated as a dependent variable (Figure 2). Note that we applied a natural logarithmic transformation to the averaged centroid data to fulfill the assumption of normal distribution and tested this using the Shapiro–Wilk test.

Another interest in the study is whether the index of brain network flexibility serves as a proxy for the trial-by-trial variability of musical performance. First, each independent and dependent variable used for this regression analysis was standardized at the individual level as follows: z_*i*_=(x_*i*_-μ)/σ. Second, the normalized trial-by-trial individuals’ data were serially concatenated. Lastly, this shaped dataset was fed into the elastic net regression. Elastic net regression was performed using the *glmnet* package implemented into R (Friedman et al. 2010). The goodness-of-fit of the models was verified by using a coefficient of determination value (R^2^). Statistical significances were tested at p < 0.05.

## Results

Note that 33.7 % of the trials across all participants were excluded from our reports because of excessive noise in the preprocessed EEG signals or issues with musical performance.

The Shapiro–Wilk test revealed that the log-transformed centroid data followed a normal distribution (p>0.05). The group-averaged index of brain network flexibility in each frequency band at each cortical region of interest is depicted in Figure 3. The two-way repeated measures ANOVA found a significant main effect of Frequency bands (F_2.4,69.2_=13.9, p<0.0001) and Brain regions (F_1.3,36.3_=197.9, p<0.0001), but no interaction between them (F_3.6,101.4_=1.19, p=0.31). The post-hoc test found that brain network flexibility significantly differed among the brain regions. Specifically, the occipital brain region had the most flexible network of the predefined four brain regions (p < 0.001, adjusted for multiple comparisons). With respect to the effect of frequency bands, the theta frequency band had significantly higher brain network flexibility (p < 0.005, adjusted for multiple comparisons) than the other frequency bands, indicating that the theta frequency had the most flexible network (Figure 3B). Based on these behavioral and neural features, we aimed to clarify the extent to which brain network flexibility is associated with musical performance in expert musicians at the between-subject level. To address this, we developed a statistical model using elastic net regression associating musical performance (i.e. sound centroid) with brain network flexibility. Figure 4A summarizes the results of elastic net regression. Our model selected the frontal region in the theta and beta1, the parietal region in the alpha and beta1, the temporal region in the theta and beta1, and the occipital region through theta to beta2 frequency as key dependent variables that were highly associated with skillful musical performance. The coefficient of determination value between the observed and predicted values of the Centroid was 0.80 (Figure 4B).

**Figure 4:**
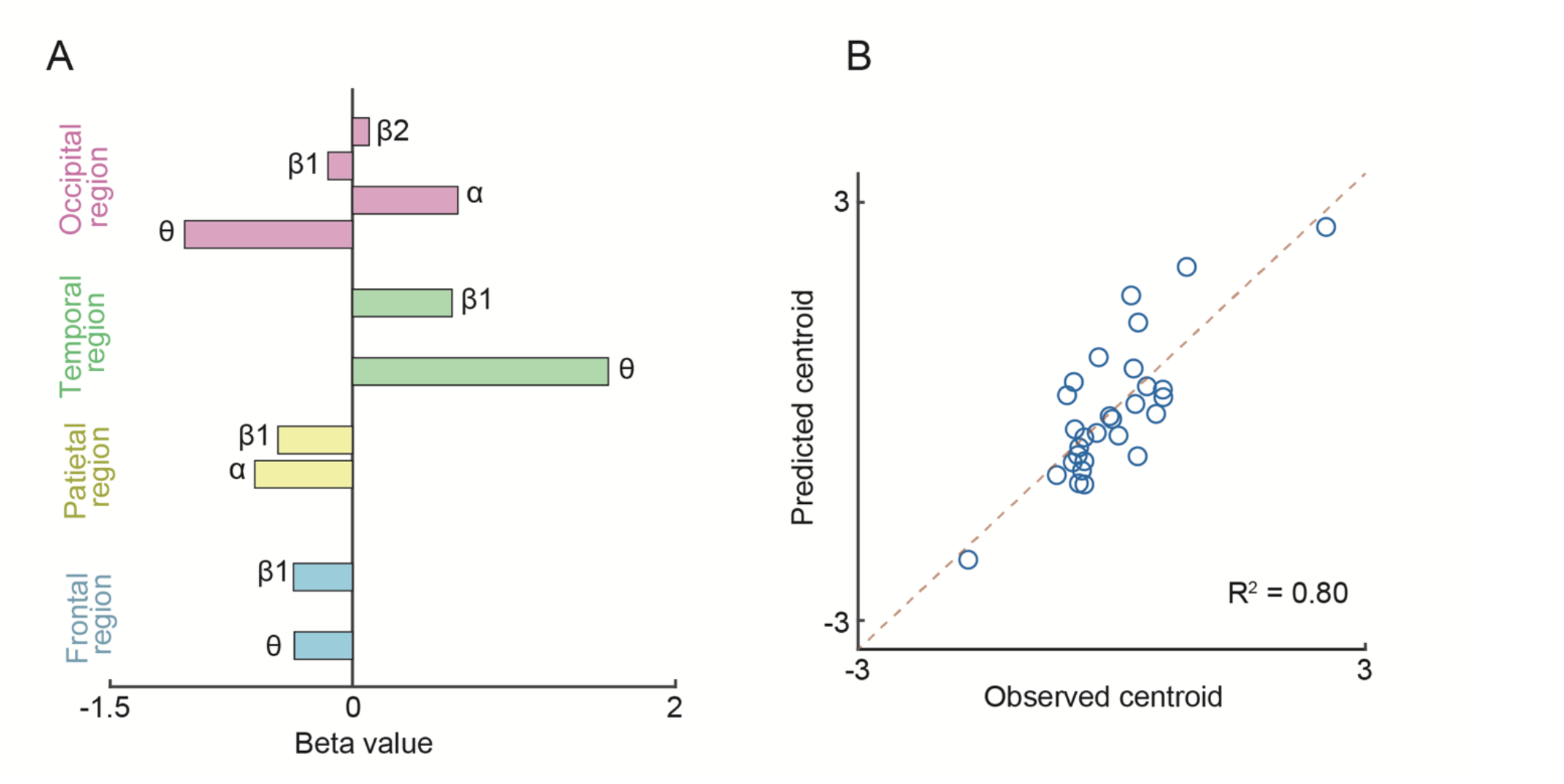
The result of elastic net regression. (A) Each bar plot shows beta values computed by the elastic net regression. This regression revealed that widespread brain regions and frequency bands were tightly associated with skilled musical performance. (B) Observed versus predicted centroid values from the model are shown. Each dot represents an individual. The R^2^ value of this regression was 0.80.

Furthermore, we investigated whether brain network flexibility is capable of predicting the trial-by-trial variability of musical performance. However, no reliable regression model was found in our dataset (R^2^ was less than 0.1). Thus, predicting single-trial musical performance from brain network flexibility was almost impossible in this case.

## Discussion

By combining EEG recordings, behavioral measurements of musical performance, and mathematical approaches, we aimed to determine whether brain network flexibility prior to musical performance can predict inter-individual and intra-individual variability in skilled musical performance among brass instrumentalists. Our results suggest that brain network flexibility can predict inter-individual differences in precision of tone timbre (as represented by spectral centroid of the sound), but not trial-by-trial variability at the individual level. Furthermore, we attempted to identify whether regional differences in brain network flexibility are related to skilled musical performance. We found that brain network flexibility in broader cortical regions, rather than specific local cortical regions, predicted skilled musical performance. Thus, we propose that flexible large-scale brain networks may potentially serve as a proxy for inter-individual differences in skilled musical performance.

In the past decade, human fMRI studies have reported BOLD-based brain network flexibility during the period of 5- to 10-mins of a resting/task-free state (Smith et al. 2009; van den Heuvel and Pol 2010; Smith Stephen M et al. 2013) and its relation to memory, learning, cognition, and motor functions in humans. However, this network flexibility was obtained in the absence of a task or experimental stimuli, and task-irrelevant brain network flexibility was thus used to predict motor and cognitive function. One reason for this is that the temporal resolution of fMRI is constrained by hemodynamic response time, which peaks around 5 seconds. This indicates that the temporal information on neural activity is heavily blurred. In addition, a relatively long scan time is required to calculate brain network flexibility due to low temporal resolution. Therefore, the original fMRI-based network flexibility (i.e., resting-state fMRI) seems inadequate for determining the brain’s response to online task demands such as those immediately before and during motor execution for a single trial, which matters particularly in skillful motor behaviors such as musical performance. To deal with this methodological limitation, we leveraged multi-channel scalp EEG recordings that have an excellent temporal resolution in a millisecond order. Hence, EEG-based brain network flexibility can determine the high temporal resolution of brain network flexibility even under a narrow time window during the motor preparation. Our approach therefore offers a complement to pioneering research that quantified brain network flexibility using BOLD signals (Bassett et al. 2011) and provides clues regarding brain network flexibility in skillful behaviors that prioritize more temporal information.

In this study, we found a significant difference in that brain network flexibility among cortical regions and frequency bands, with the frontal region being less flexible compared to other regions. Two possible interpretations of this finding are suggested. First, the frontal region has been regarded as a key functional hub of the brain networks (Power et al. 2013; Marek and Dosenbach 2018) and may exhibit unique activation patterns compared to other regions, as it is known to orchestrate multiple brain networks during a task (Cole et al. 2013). Second, the “task buffer” theory (Cole et al. 2017) proposes that the frontal region acts as a buffer for maintaining multiple task representations without interference with ongoing task performance. During the preparation period in this study, participants had to prepare for both upcoming Go and NoGo trials in parallel, and we suggest that altered frontal network flexibility may serve this “task buffer” function.

The most important finding in the present study was that brain network flexibility throughout the entire cortical regions was involved in predication of skilled musical performance. Specifically, the frontal region in the theta and beta1 (13-25Hz) frequency bands, the parietal region in the alpha and beta1 frequency bands, the temporal region in the theta and beta1 frequency bands, and the occipital region through theta to beta2 (26-40Hz) frequency bands were the most influential explanatory factors. This finding indicates that whole-cortical network flexibility, rather than network flexibility in a specific local cortical region, may determine skilled musical performance. Interestingly, broader EEG frequency ranges from theta to high beta bands were also involved. This suggests that whole-cortical dynamics over time and cortical regional space are inextricably associated with individual musical performance. Whole-brain modes, as proposed by Deco ang colleagues (Deco et al. 2009), provides a key idea that flexible communication between brain regions is necessary for the optimal exploration of the effective dynamical repertoire of brain networks. If such optimal exploration is much more limited at the mesoscopic level for some reason (e.g., dysfunction), individuals cannot flexibly update their behavioral strategies during musical performance. Viewed in this light, our results suggest that individuals with skilled musical performances possess effective dynamical repertoires of brain networks.

In this study, we are further interested in whether brain network flexibility can predict not only inter-individual variation but also trial-by-trial variability in skilled musical performance. This is because neural activity is known to fluctuate over time even when individuals receive repeated presentation of the same stimulus (Arieli et al. 1996; Faisal et al. 2008), and these fluctuations may cause considerable variability of motor skill across trials. Furthermore, skilled musical performance at each trial may be influenced by brain network flexibility resulting from fluctuations in neural activity. However, contrary to our hypothesis, the present study did not show strong relationships between trial-by-trial variability of the performance and brain network flexibility. One possible explanation is the lower signal-to-noise ratio of a single trial of the brain’s electrical signal. The electrical signals in the brain induced by the stimulus or response are assumed to be small because they are mixed with continuous background activity irrelevant to the given stimulus or response (Blythe LaGasse et al. 2022). Based on our results, we suggest that the averaged brain network flexibility over trials is useful for predicting inter-individual differences in skilled motor performance, but not for explaining trial-by-trial variability in individual performance.

To conclude, we show here, for the first time, that EEG-based brain network flexibility is capable of predicting skilled musical performance in expert brass instrumentalists. We found that EEG-based regional brain network flexibility was tightly associated with inter-individual differences in musical performance, but not with trial-by-trial variability within the individual level. Although further studies are needed, such as a causal approach with non-invasive brain stimulation, to provide solid evidence supporting our findings, our novel findings have significant potential for designing a new approach to predict an individual’s skill from neural dynamics and a new intervention tool, such as a neurofeedback system, to facilitate musical training.

## Acknowledgements

We thank to all participants for taking part in this study. This work was supported by Japan Science and Technology Agency, Moonshot R&D Grant Number JP-MJMS2012

## Figure legends

Figure 1: Experimental setup and trial structure. (A) Participants sat upright in a chair in front of a monitor while holding their instruments with both hands. After each trial, they used a foot pedal mouse controlled by their right foot to self-evaluate the quality of their musical performance on a visual analog scale. Acoustic signals of the long-tone sound were recorded using a microphone in a silent environment. Throughout the data collection, 32-channel scalp EEG signals were continuously recorded. (B) Participants instructed to perform a visually-guided Go/Nogo task with playing a long-tone using their own musical instrument. On every trial, participants prepared to simultaneously either play a long-tone or cancel musical performance. After 5 seconds of rest and receiving around 2 seconds of the ready cue (yellow circle), one of two different colors of cues was presented in random order within the session. If participants received the visual “Go” cue (green circle), they were asked to play a long tone for 5 seconds as accurately as possible. On the other hand, if they received the visual “NoGo” cue (red circle), they inhibit their musical performance as quickly as possible. At the end of each trial, participants rated their performance on a visual rating scale from 1 (worse) to 7 (better) using the foot pedal mouse connected to a PC.

Figure 2: Pipeline for analyzing acoustic and neural data. The spectral centroid, which represents the brightness of timbre while playing a long-tone, was calculated using a spectral-based analysis. The variability of the spectral centroid obtained during a 3 second window was used as a dependent variable. For the neural feature, we first computed dynamic PSI matrices using the phase difference between all channel pairs. The data from these dynamic PSI matrices were fed into a community detection analysis using the Louvain algorithm. The indices of brain network flexibility in four predefined brain regions (frontal, temporal, parietal, and occipital regions) and four frequency bands (theta, alpha, beta1, and beta2) were treated as independent variables. To reveal the relationship between behavioral and neural features, a model of Elastic Net regression was used.

Figure 3: An anatomically-based parcellation of the EEG channels and results of brain network flexibility across each brain region and frequency band during the preparation period. (A) Thirty-two channels, excluding the two mastoid channels, were classified based on the neuroanatomical locations. Four predefined brain regions of interest were classified: the frontal region (8 channels: Fp1, Fp2, F3, F4, Fz, F7, and F8), the temporal region (6 channels: T7, T8, CP5, CP6, P7, and P8), the parietal region (12 channels: FC1, FC2, FC5, FC6, C3, C4, Cz, CP1, CP2, P3, Pz, and P4), and the occipital region (4 channels: POz, O1, O2, and Oz). (B) Each grey bar plot shows the group-averaged flexibility indices for each brain region and frequency band. The y-axis represents the level of flexibility in brain networks, with high values indicating greater flexibility. Each dot represents an individual, and each error bar indicates the standard error of means across individuals. Three asterisks indicate p < 0.0001.

Figure 4: The result of the elastic net regression. (A) Each bar plot shows beta values computed by the elastic net regression. This regression revealed that widespread cortical regions and frequency bands were tightly associated with skilled musical performance. (B) Observed versus predicted centroid values from the model are shown. Each dot represents a datapoint of each individual. The R^2^ value of this regression was 0.80.

